# Mesenchymal stem cell subpopulations and their heterogeneity of response to inductions revealed by single-cell RNA-seq

**DOI:** 10.1101/2021.05.07.443197

**Authors:** Wenhong Hou, Li Duan, Changyuan Huang, Xingfu Li, Xiao Xu, Pengfei Qin, Ni Hong, Daping Wang, Wenfei Jin

## Abstract

Mesenchymal stem/stromal cells (MSCs) are promising cell source for regenerative medicine and treatment of autoimmune disorders. Comparing MSCs from different tissues at single cell level is fundamental for optimizing clinical applications. Here we analyzed single cell RNA-seq data of MSCs from 4 tissues, namely umbilical cord, bone marrow, synovial tissue and adipose tissue. We identified 3 major cell subpopulations, namely osteo-MSCs, chondro-MSCs, adipo/myo-MSCs, across all MSC samples. MSCs from umbilical cord exhibited the highest immunosuppression, potentially indicating it is the best immune modulator for autoimmune diseases. The differentiation potentials of MSC subpopulations, which are strongly associated with their subtypes and tissue sources, showed pronounced subpopulation differences. We found MSC subpopulations expanded and differentiated when their subtypes consist with induction directions, while the other subpopulations shrank. We identified the genes and transcription factors underlying each induction at single cell level and subpopulation level, providing better targets for improving induction efficiency.

## Introduction

Mesenchymal stem/stromal cells (MSCs) are multipotent stromal cells that can differentiate into a variety of cell types including osteoblast, chondroblast, osteocyte, myocyte and adipocyte *in vitro* [1–4]. The promising features of MSCs, including self-renewal capacity and ability to differentiate into different cell types, have aroused great interests among scientists and medical experts whose work provided an attractive perspective on MSCs-based tissue repairs [3, 5–7]. Further studies showed MSCs also played an important role in tissue homeostasis and immunomodulation via interaction with immune cells and secretion of various factors including growth factors, cytokines and antifibrotics [5, 6, 8–10]. MSCs have become the most promising cell source for cell-based therapies, particularly in tissue repair and treatment of immune disorders [5, 7, 11–13]. The number of clinical trial on MSCs-based therapies is rising considerably in recent years, indicating the great potential of MSCs. However, dysregulation of MSCs induction and low efficiency of MSCs induction into target functional cells remain hinders the application of MSC-based therapies [14–17].

Although there are established approaches for induction of MSCs into specific functional cell type *in vitro* [14, 18–20], Most studies showed that MSCs exhibited significant differences in colony morphologies, proliferation rates and differentiation potentials [21–23]. The lack of clear understanding of cellular heterogeneity of MSCs severely hampered the development of an efficient and reproducible clinical application [24, 25]. Furthermore, MSCs have been identified in most tissues in our body and are isolated from bone marrow, umbilical cord, adipose tissue, synovial tissue, muscle, liver, dental pulp and so on [26–30]. It remains largely unexplored whether MSC subpopulations are consistent across tissues. In particular, the response heterogeneities of MSCs from different tissue sources to inductions are unknown. Furthermore, our knowledge on lineage commitment of MSCs are mainly based on analysis of bulk assay, which captures the average signal across entire populations while ignored response heterogeneities of MSCs.

Single cell RNA-seq (scRNA-seq) is very powerful to reveal cellular heterogeneities under various conditions [31–37]. Here we analyzed scRNA-seq data of MSCs from multiple tissues to elucidate MSC subpopulation across tissues. We found the MSC subpopulations from different tissues were essentially consistent with each other, although the abundance of each subpopulation was highly diverse across tissues. We further analyzed the processes of MSCs induction into different functional cells to provide novel insight on mechanisms of MSCs differentiation. We further identified transcription factors underlying each induction at single cell level to elucidate the potential target for efficient induction.

## Results

### scRNA-seq showed consistent MSC subpopulations across tissues

Human umbilical cords (UC) were collected from Shenzhen Second People’s Hospital, with donors signed informed consent approved by the IRB of the hospital. UC-MSCs were isolated from UC Wharton’s jelly following Reppel *et al.* [38] (see Methods). Two UC-MSC samples and a synovial tissue (SY)-MSC sample were used for generating scRNA-seq data following 10X genomics protocol [39]. Majority of the cells from the two UC-MSCs samples expressed *THY1*, *NT5E* and *ENG* (Fig. S1a). The scRNA-seq data of MSCs from bone marrow (BM) [40] and adipocytes (AD) [41] were integrated for analyses of the cellular heterogeneities of MSCs across different tissues. All the cell subpopulations from the 4 tissues expressed MSCs specific markers such as *THY1*, *NT5E* and *ENG* (Fig. S1b). Unsupervised clustering each of the 4 MSCs samples resulted in 3 distinct clusters, respectively (Fig. S1c, S1e, S2a, S2c). We still identified 3 major cell subpopulations on UMAP projection after we integrated the 4 MSC samples (Fig. 1a). Each MSC sample has the 3 subpopulations but with different abundances (Fig. 1b), indicating the major MSC subpopulations are consistent among these MSC samples. The three MSC subpopulations exhibited lineage specific expression, namely chondrocyte lineage (*HMGB1*, *HMGB2, DCN, F3*, *MDK, BMP5, KIAA0101*), adipocyte/myocyte lineage (*FTL*, *FTH1*, *TAGLN, FKBP1A*, *ACTG2*, *TXN,*) and osteoblast lineage (*BGN*, *HAPLN1*, *FHL1*, *VCAN*, *GDF15*) (Fig. 1c, S1a, S1f, S2b, S2d). We named the three subpopulations as chondrocyte lineage MSCs (chondro), adipocyte/myocyte lineage MSCs (adipo) and osteoblast lineage MSCs (osteo). MSCs deriving from different tissues while clustering into the same subpopulation exhibited similar lineage specific expression profiles (Fig. 1c, 1d), further indicating all the 4 MSC samples had the same cell subpopulations.

**Figure 1.**
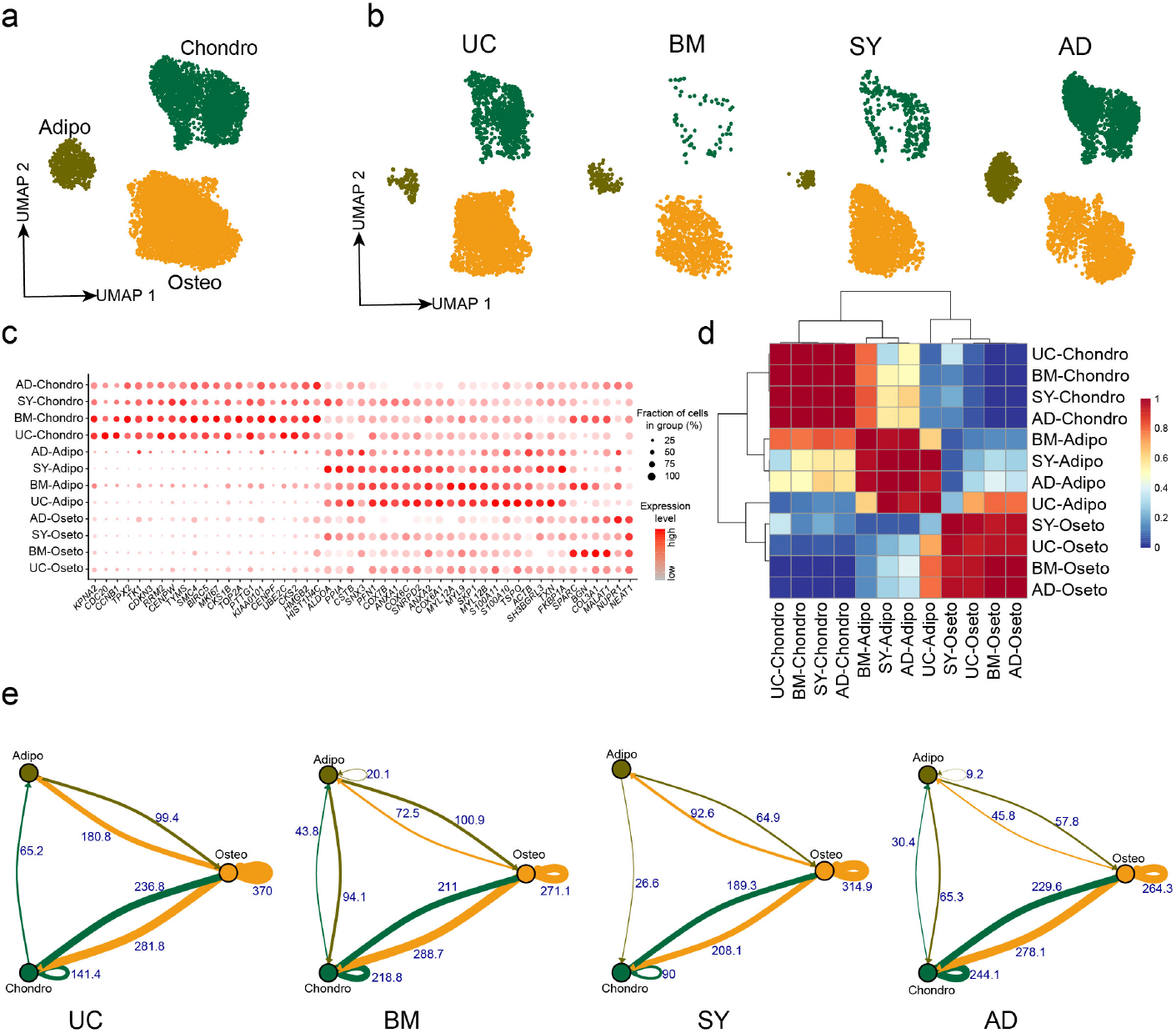
MSCs from different tissues have similar cell subpopulations. a. UMAP projection of MSCs from UC, BM, SY and AD. b. UMAP projection of MSCs from each sample, namely UC-MSCs, BM-MSCs, SY-MSCs and AD-MSCs. c. Dot plot showing the expressions of 3 MSC subpopulation specific genes in 4 tissues. d. AUROC score across cell subpopulations from the 4 tissues (UC-MSCs, BM-MSCs, SY-MSCs and AD-MSCs) by MetaNeighbour. e. Cell-Cell crosstalk among the three cell subpopulations in MSCs from each tissue, namely UC-MSCs, BM-MSCs, SY-MSCs and AD-MSCs.

The crosstalk of ligand–receptor on cell surface play an important role in cellular signaling transduction. We analyzed the crosstalk of ligand–receptor pairs to understand the cell-cell communication in MSCs. We found osteo→osteo and osteo→chondro showed the strongest interaction among all subpopulation pairs in each MSC sample (Fig. 1e, S2e-h). Therefore, oseto is the most important signaling sender in MSCs, potentially indicating osteo shape and contribute the cellular micro-environment. The crosstalk between osteo and chondro is the second strongest, potentially indicating there are frequent cell-cell communication between them. The crosstalk between adipo and osteo/chondro is very weak, potentially indicating that adipo are relative isolated in MSCs. The observations are consistent with recent reports that adipogenesis and osteogenesis/chondrogenesis are mutually repressive processes [4, 40].

### MSC subpopulations from different tissues show different potentials

It is essential to compare the differentiation potentials and immunomodulatory ability of MSC subpopulations from different tissue sources due to their implications in MSC-based therapy. UC-MSCs always exhibited the highest stemness/entropy among all MSC samples (Fig. 2a), potentially indicating their strongest proliferation potential and strongest stemness. Indeed, stemness-associated genes including *AMIGO2*, *CLDN1*, *LRRC17* and *SLC22A3* were significantly highly expressed in UC-MSCs (Fig. 2b). Furthermore, violin plot showed UC-MSCs have the highest immunosuppression scores among all MSC samples (Fig. 2c), e.g. *AREG*, *CSF3*, *CCL20* and *IL6* were significantly highly expressed in UC-MSCs (Fig. 2d). These results potentially indicate that UC-MSCs are the best MSC source for immunomodulation of innate and adaptive immune responses, consistent with the reports using bulk data [42].

**Figure 2.**
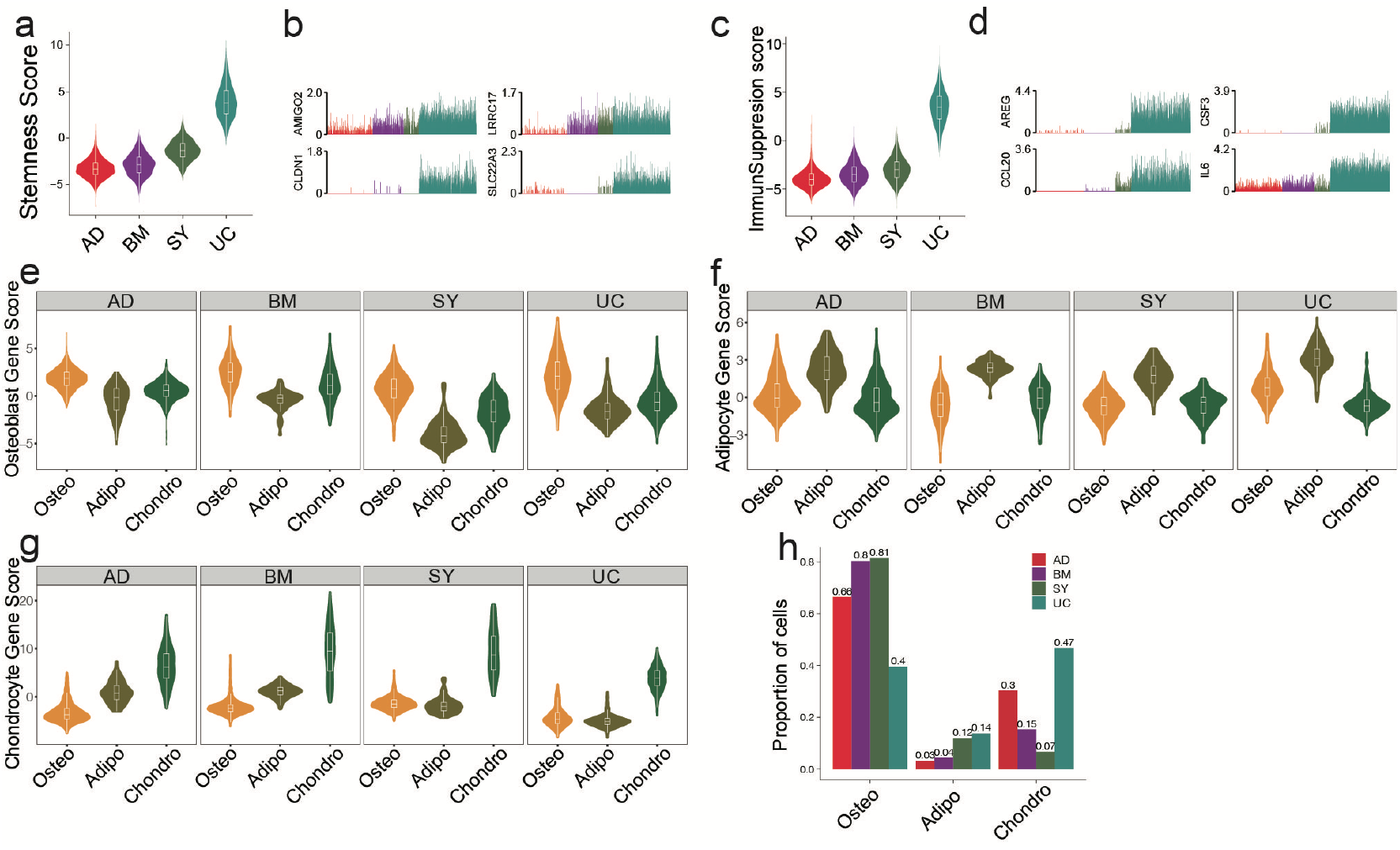
MSC subpopulations from different tissues have different potencies. a. Violin plot showing stemness scores of MSCs from 4 different tissues. b. Bar plot showing expression of *AMIGO2, CLDN1, LRRC17* and *SLC22A3* in each cell of the MSCs. c. Immunosuppression score of MSCs from 4 tissues. d. Bar plot showing expression of four immunosuppression genes *AREG, CSF3, CCL20* and *IL6* in each gene. e-g. Osteoblast score (e), adipocyte score (f) and chondrocyte score (g) of MSC subpopulations from the 4 tissues. h. Abundances of MSC subpopulations in different tissues.

Investigation of differentiation potential of MSCs to osteoblast lineage, chondrocyte lineage and adipocyte/myocyte lineage are also crucial for clinical applications. The osteo subpopulations from the 4 tissues exhibited the highest differentiation potential to osteoblast, among which BM-osteo and UC-osteo had the highest osteoblast gene scores (Fig. 2e). On the other hand, SY-adipo exhibited the lowest differentiation potential to osteoblast, indicated it had to overcome much high barriers to differentiate to osteoblast (Fig. 2e). Similar analyses showed that AD-adipo and BM-chondro had the highest differential potential to adipocyte lineage and to chondrocyte lineage, respectively (Fig. 2f, 2g). Analyses of the abundances of MSC subpopulations from different tissues showed BM and SY have the highest fractions of osteo, potentially because both tissues are osteo associated (Fig. 2h).

### Heterogeneous response of MSCs to chondrogenesis induction

Induction of MSCs to chondrocyte-like cells have important implications for cartilage and bone regeneration [3]. Since UC-MSCs have the highest fraction of chondro, UC-MSCs was induced to chondrocyte-like cells following our previous study [43]. We conducted scRNA-seq on chondrogenesis-induced MSCs and obtained 6,161 high quality single cell transcriptomes for further analyses. We merged pre-induced MSCs and chondrogenic-induced MSCs for better understanding of their relationships (Fig. 3a). We found chondrogenic-induced osteo and chondrogenic-induced adipo partially overlapped with their counterparts in pre-induced UC-MSCs on t-SNE plot (Fig. 3b, S3a). On the other hand, chondrogenic-induced chondro and its counterpart in pre-induced MSCs are complete separated from each other (Fig. 3b), indicating the states of chondro changed a lot after chondrogenesis induction. The expressiones of lineage specific genes, such as *HMGB2*, *HIST1H4C*, *BGN*, *S100A10*, *TXN*, *FKBP1A*, *CXCL2*, *IL6, FGF2, MDK* and *MMP14*, are essentially consistent pre-and post-induction, although their expression varied somewhat (Fig. 3c, S3c). Chondrogenic-induced chondro has significant higher chondrocyte score compared to pre-induced chondro (Fig. 3d), indicating the chondrogenesis induction works well. The fraction of chondro in chondrogenic-induced MSCs (91%) is much higher than its counterpart in pre-induced MSCs (67%) (Fig. 3e), indicating chondro expansion during the induction. On the other hand, the fractions of oseto (1.6%) and adipo (7.7%) in chondrogenic-induced MSCs were much lowers than that of their counterparts in pre-induced MSCs (Fig. 3e), thus both oseto and adipo relatively shrank, potentially due to the induction are unfavorable for their proliferation. Entropies of chondrogenic-induced MSC subpopulations were significantly lower than their counterparts in pre-induced MSCs (Fig. S3b), implying the decreased stemness during induction. These results indicated both dynamics of cell subpopulation and changes of cell status played roles during chondrogenesis induction.

**Figure 3.**
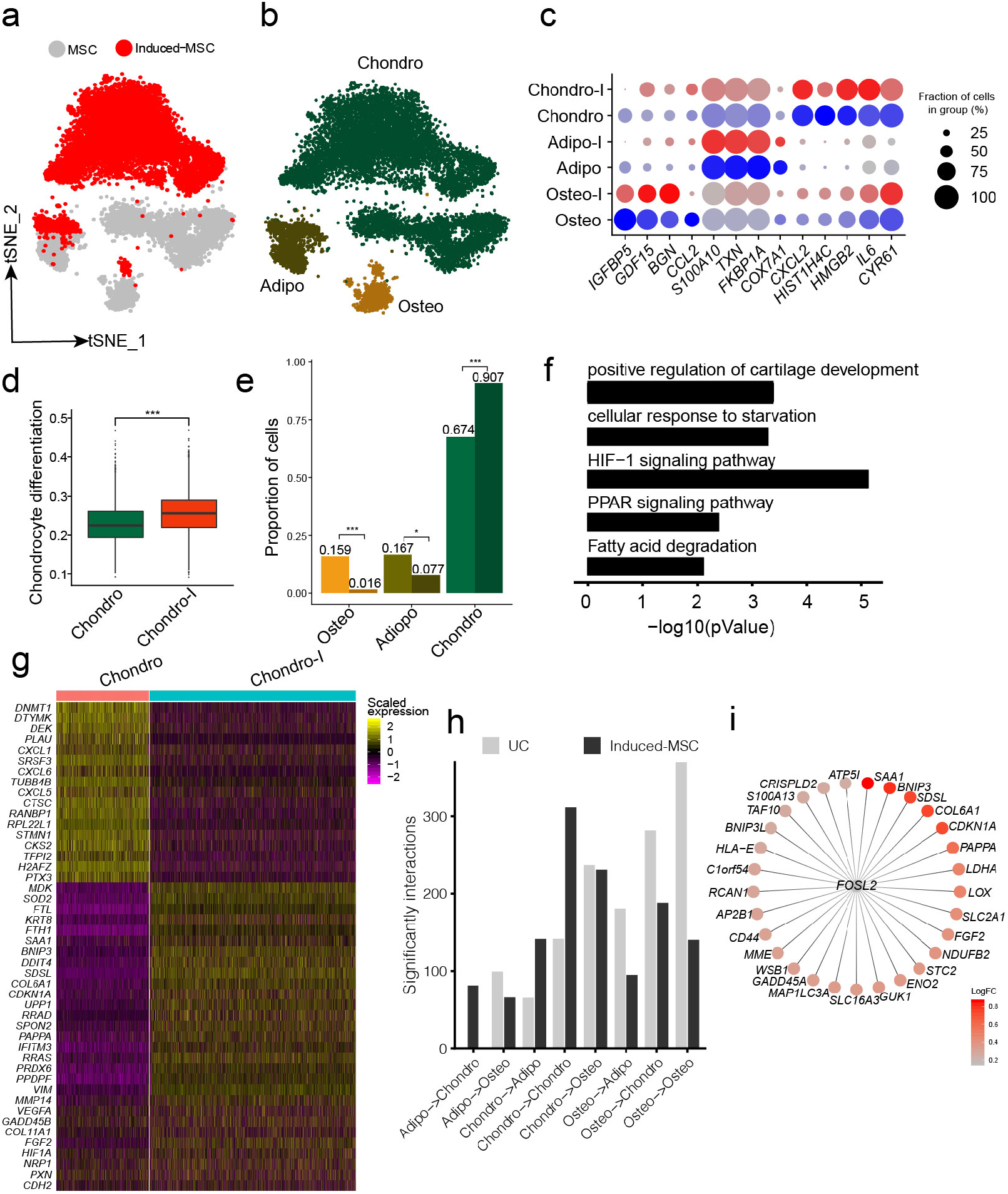
Heterogeneous response of MSCs to chondrogenesis induction. a. t-SNE projection of UC-MSCs and chondrogenic-induced MSCs, colored by UC-MSCs and chondrogenic-induced MSCs. b. t-SNE projection of UC-MSCs and chondrogenic-induced MSCs, colored by MSC subpopulation, namely chondro, osteo and adipo. c. Consistent expression of lineage specific genes in MSCs subpopulations pre and post induction. d. Boxplot of chondrocyte differential scores of chondro and and chondrogenic-induced chondro. e. Fractions of MSC subpopulation in MSCs and chondrogenic-induced MSCs. * stands for p value smaller than 0.01, *** stands for p value smaller than 0.0001. f. GO analysis of up-regulated genes in chondrogenic-induced chondro compared to pre-induced chondro. g. Heatmap of differentially expressed genes between pro-and post-induced chondro. h. Chondro-chondro crosstalk is significantly increased after chondrogenesis induction. i. *FOSL2* and its target genes colored by logFC of chondro between MSCs and chondrogenic-induced MSCs.

We identified 393 significantly differentially expressed genes between pre-induced chondro and chondrogenic-induced chondro. The significantly up-regulated genes in chondrogenic-induced chondro are enriched in positive regulation of cartilage development (4.6×10^−4^), HIF-1 signaling pathway (7.9×10^−6^), PPAR signaling (4.5×10^−3^) and so on (Fig. 3f). The most up-regulated genes include *COL6A1*, *COL6A2*, *DCN*, *FGF2*, and *MMP14* (Fig. 3g), which play important roles in collagen formation or priming chondrogenic progenitors [44, 45]. The up-regulated genes include hypoxia-associated genes such as *HIF1A*, consistent with previous report that low oxygen level could promote chondrogenic differentiation [46]. The most down-regulated genes in chondrogenic-induced chondro were cell cycle-related genes (*BIRC5*, *CCND1*, *PCNA*, *TPX2*, *AURKA*), consistent with aforementioned reduced stemness and more differentiated states. On the other hand, the osteoblast lineage specific genes (*COL1A1*, *COL1A2, SPARC*, *TGFBI*) and adipocyte/myocyte lineage specific genes (*FKBP1A*, *TAGLN*) have been down-regulated in induced osteo and induced adipo, respectively (Fig. S3d, S3e). These observations indicate chondrogenesis induction promotes chondro differentiation while represses adipo differentiation and oseto differentiation, indicating the highly heterogeneous response of MSCs.

The cell-cell crosstalk of chondro-chondro significantly increase after chondrogensis induction (Fig. 3h), which may indicate the crosstalk between chondros played an important role during chondrogensis induction. In order to better understand the processes and mechanisms during chondrogenesis induction, we identified induction associated transcription factors (TFs) networks. The top up-regulated TFs networks are *FOSL2, ATF5, FOXF1, HES7*, and so on, among which *FOSL2* regulated a lot target genes include *LDHA*, *SAA1*, *BNIP3*, *COL6A1*, *CDKN1A*, *FGF2* (Fig. 3i) [47–49], potentially indicating *FOSL2* plays a key role in chondrogenesis induction.

### Heterogeneous response of MSCs to osteogenesis induction

Our comparison analyses showed BM-MSCs have the highest fractions of osteo and osteo from BM-MSCs showed the highest differentiation potential to osteoblast among all MSC samples. Therefore, BM-MSC is the best candidate for induction of MSCs to osteoblast. scRNA-seq data of BM-MSCs and osteogenic-induced BM-MSCs were obtained from Rauch *et al.* [40]. The BM-MSCs and osteogenic-induced BM-MSCs were essentially overlapped on integrated t-SNE projection (Fig. 4a). The MSCs were clustered into oseto, chondro and adipo (Fig. 4b, S4c, S4d), same as aforementioned. The expressions of lineage specific genes, such as *HMGB2*, *PDGFRA, TMEM119*, *HIST1H4C*, *ACTB, TAGLN, IL6, FGF2* and *MMP14*, are essentially consistent pre- and post-induction, although their expression varied somewhat (Fig. 4c). The osteogenic-induced osteo exhibited enhanced osteoblast proliferation compared to pre-induced osteo (Fig. 4d), indicating the osteogenesis induction works well. The fractions of osteo and chondro increased little in osteogenic-induced MSCs, while the fraction of adipo significantly decreased (Fig. 4e), consistent with the report that osteogensis and chondrogensis sharing part of the development trajectory while osteogensis and adipogensis being opposite process [4, 40]. The osteogenic-induced MSC subpopulations had lower entropies than their counterparts in pre-induced MSCs (Fig. S4a), consistent with the observation that differentiated cells have lower entropies/stemnesses.

**Figure 4.**
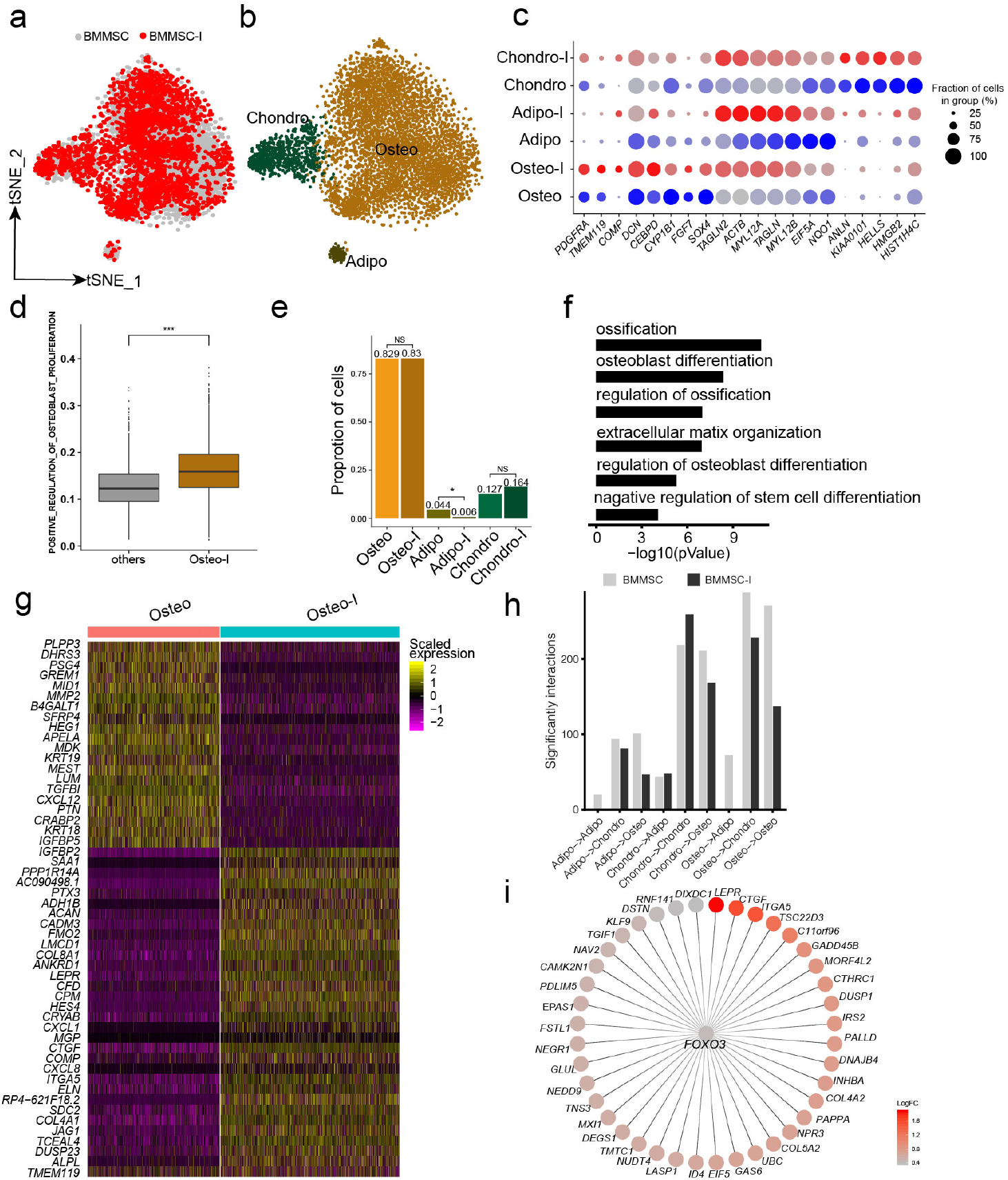
Heterogeneous response of MSCs to osteogenesis induction. a. t-SNE projection of MSCs and osteogenic-induced MSCs, colored by MSCs and osteogenic-induced MSCs. b. t-SNE projection of MSCs and osteogenic-induced MSCs, colored by MSC subpopulation, namely chondro, osteo and adipo. c. Consistent expression of lineage specific genes in MSC subpopulations pre and post osteogenesis induction. d. Boxplot of osteoblast differential scores of chondro pre and post induction. e. Change of cell subpopulations size pre and post osteogenesis induction. * stands for p value smaller than 0.01, *** stands for p value smaller than 0.0001. f. GO analysis of up-regulated genes in osteogenic-induced osteo. g. Heatmap of differentially expressed genes between pro-and post-induced osteo. h. Cell-Cell crosstalk among the three MSC subpopulations. i. *FOXO3* and its target genes colored by logFC of osteo between MSCs and osteogenic-induced MSCs.

We identified 1,328 significantly differentially expressed genes between pre-induced osteo and osteogenic-induced osteo. The significantly up-regulated genes in osteogenic-induced osteo enriched in ossification (1.6×10^−11^), osteoblast differentiation (3.2×10^−9^), extracellular matrix organization (1×10^−7^), regulation of osteoblast differentiation (1×10^−7^) and negative regulation of stem cell differentiation (6.3×10^−5^) (Fig. 4f). The most up-regulated genes include *ACAN*, *COL8A1*, *HES4*, *CTGF*, *ITGA5*, *JAG1*, *ALPL*, *TMEM119* and *COMP* (Fig. 4g), among which *ALPL*, *COMP* and *TMEM119* are well-known osteoblast specific genes (Fig. S4b). Interestingly, the cell-cell crosstalk of osteo-osteo didn’t show increase after osteogensis induction (Fig. 4h), but with strong cell-cell crosstalk with chondro, which may indicate the crosstalk between the chondro and osteo may play important role in osteogensis induction. We further identified osteogensis associated TF networks such as *MAFF, FOXO1, MXI1*, among which *FOXO3* up-regulated a lot gene including *LEPR*, *CTGF*, *ITGA5* and *COL5A2* (Fig. 4i), consistent with recent studies [50, 51].

### Heterogeneous response of MSCs to adipogenesis induction

We further analyzed the responses of BM-MSCs subpopulations to adipogenesis using scRNA-seq data from Rauch *et al.* [40]. Unsupervised clustering of MSCs from both BM-MSCs and adipogenic-induced BM-MSCs identified 4 MSC subpopulations, namely oseto, chondor, adipo and myoblast (myo) (Fig. 5a, 5b, S5a, S5d, S5e, S5f). Adipo emerged after adipogenesis induction since there is almost no adipo in pre-inducted MSCs, while the fraction of the other 4 subpopulations decreased (Fig. 5c). All subpopulations in adipogenic-induced MSCs had lower entropies compared with their counterparts in pre-induced MSCs, indicating a more differentiated cell state after adipogenesis induction (Fig. S5c).

**Figure 5.**
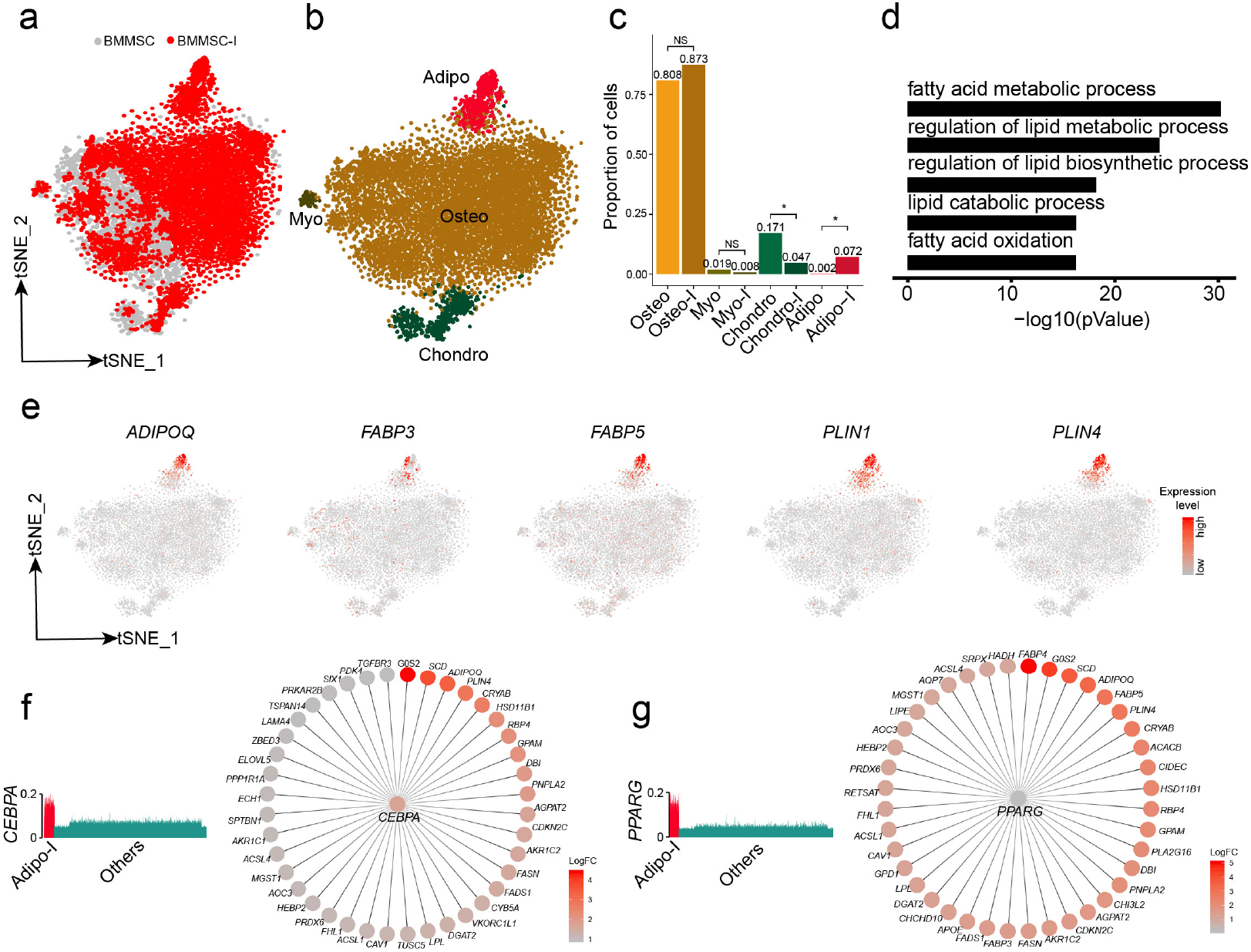
Heterogeneous response of MSCs to adipogenesis induction. a. t-SNE projection of MSCs and adipogenic-induced MSCs, colored by MSCs and adipogenic-induced MSCs. b. t-SNE projection of MSCs and adipogenic-induced MSCs, colored by chondro, osteo, adipo and myo. c. Change of fractions of MSC subpopulations pre and post adipogenesis induction. *stands for p value smaller than 0.01, *** stands for p value smaller than 0.0001. d. GO analysis of induced adipo. e. Expression of typical adipogenic lineage specific genes on t-SNE projection of MSCs f. *CEBPA* and its target genes colored by logFC of adipo between MSCs and adipogenic-induced MSCs.

Adipogenic-induced adipo significantly enriched genes associated with adipocytes, such as fatty acid metabolic process (8×10^−31^), regulation of lipid metabolic process (6.3×10^−25^) and regulation of lipid biosynthetic process (6.3×10^−19^) (Fig. 5d). The adipocyte lineage specific genes, such as *ADIPOQ*, *FABP3*, *FABP5*, *PLIN1* and *PLIN4* [52–54], almost exclusively expressed in adipogenic-induced adipo (Fig. 5e). We identified the TF networks, such as *CEBPA, PPARG* and *STAT5A*, played important role during adipogenesis induction. *CEBPA* and *PPARG* networks co-regulated a lot adipogenic associated genes such as *ADIPOQ*, *PLIN4*, *FABP4* and *FABP5* (Fig. 5f, 5g), consistent with previous studies [55, 56]. The results indicated that adipogensis induction lead quite significant transcriptome change, different from the mild change during chondrogensis induction and osteogensis induction.

## Discussion

It is well known that MSCs exhibit considerable tissue-to-tissue heterogeneity and intra-subpopulation heterogeneity [1]. However, we have very limited knowledge about whether MSC subpopulations across different tissues are consistent and heterogeneous responses of MSCs from different tissues to the lineage specific inductions. Here, scRNA-seq analyses showed that MSCs from different tissues had similar cell subpopulation composition, mainly composed by three major MSC subpopulations (chondro, osteo and adipo). The observations are quite different from a recent report by Huang *et al.* [57] showing very limited heterogeneity in UC-MSCs. The observation of limited heterogeneity in Huang *et al.* could attribute to very small number of cells in their study. A recent study investigated the differences in gene expression profiles between BM-MSCs and UC-MSCs[58], but missed the comparison among the their subpopulation counterparts. We directly compared the lineage specific differential potential among the subpopulation counterparts thanks to consistent MSC subpopulations across different tissues, which facilitated choosing suitable MSCs for clinical usage.

The MSC subpopulation counterparts from different tissues showed quite different differentiation potentials. We found MSC subpopulation counterparts accounted different fractions of MSCs cross tissues. scRNA-seq analyses of the MSC response to osteogenesis induction, chondrogenesis induction and adipogenesis induction showed that MSC subpopulations response to these inductions quite differently. Based on observation of osteogenesis induction and chondrogenesis induction, MSC subpopulation expanded and differentiated when they are consistent with induction inductions, while other subpopulations shrunk. Adipogenesis induction lead emergence of the adipo, thus associated significantly change of gene expression profiles on this lineage. The observations are consistent with report by Rauch *et al.* [40] that adipogenesis showed much significant transcriptomic and epigenomic changes than that of osteogenesis.

Overall, we characterized MSC subpopulations and their response heterogeneity, potentially facilitated choosing suitable MSCs for clinical usage. Further study of MSCs using single cell proteomics and single cell epigenomics could associate subpopulations with specific functions [40, 59], further facilitating the development of MSC-based clinical applications.

## Material and Methods

### *Collection and culture of* human *UC-MSCs*

The human UC were collected from Shenzhen Second People’s Hospital. All healthy voluntary donors signed informed consent approved by the IRB of Shenzhen Second People’s Hospital. UC-MSCs were isolated from Wharton jelly of umbilical cord and cultured following our previous study [43]. In brief, the UC were obtained from normal deliveries without complications throughout pregnancy according to the institutional guidelines. The UC were immediately put in physiological saline containing heparin anticoagulant and were processed within 6h after collection, storage and transportation at 4°C. The UC were cut into 3-5cm long segments under sterile environment. Vessels of umbilical cords were removed and Wharton’s jelly was collected. Wharton’s jelly was cut into small pieces (2-3 mm^3^), which were placed in petri dish with MSCs culture medium (MesenGro ®human MSC Medium, StemRD, US), 10% fetal bovine serum (FBS; Gibco, Australian) and 10μg/L basic fibroblast growth factor (bFGF; Gibco, Australian). The Wharton’s jelly blocks were cultured at 37°C in a 5% CO2 incubator. Fresh medium was added to the flasks after three days. Tissue blocks were removed after 7 days culture and the medium was replaced. Medium replacement was carried out every 72h until the cells reached an 80% confluent layer. Cells were harvested with 0.25% (w/v) trypsin plus 0.02%(w/v) EDTA (Hyclone, USA) and sub-cultured at a density of 1000 cells/cm^2^.

### Chondrogenesis induction

Chondrogenesis induction was conducted following our previous study [60]. In brief, UC-MSC was induced to chondrocyte-like cells by chondrocytes specific medium. In monolayer culture, UC-MSCs were supplemented with 0.1mM dexamethasone, 40 mg/mL L-proline, 10 μg/L transforming growth factor beta-1 (TGF-β1, Peprotech, USA), 10μg/L insulin-like growth factor-1 (IGF-1) (Peprotech, USA), and 1% insulin transferrin selenium (ITS, Invitrogen). The cells were incubated for three weeks at 37 °C in a humidified atmosphere of 5% CO_2_ and the medium changed every three days.

### scRNA-seq

The UC-MSC and chondrogenic-inducted MSC were obtained directly from the cultured cells. FACS sorting was performed on a Becton Dickinson FACSAria II (BD Biosciences, Denmark) to remove the dead cells. scRNA-seq was conducted using 10X genomics platform. Chromium Single Cell 3’Gel Bead and Library Kit (P/N 120237, 120236, 120262, 10x Genomics) were used following protocols [39]. Approximately 15,000 cells were loaded per channel. Sequencing libraries were loaded on Illumina NovaSeq 6000 with paired-end kits.

### Pre-processing of scRNA-seq data

Raw sequencing data were converted to FASTQ format with demultiplexing using Illumina bcl2fastq. Cell Ranger Single-Cell Software Suite (V2.2.0 10X Genomics; https://support.10xgenomics.com) was used to perform reads alignment, barcode demultiplexing. The reads were aligned to the hg38 reference genome. Digital gene expression matrices were preprocessed and filtered using R packages scran and scater [61]. Cells with more than 4000 expressed genes (potential doublets), less than 500 expressed genes (potential low-quality libraries), more than 10% of mitochondrial UMI counts (potential cell fragments and debris) and low expression of the housekeeping genes GAPDH and ACTB were filtered out. Normalization of UMI count was performed by first dividing UMI counts by the total UMI counts in each cell, followed by multiplication with the median of the total UMI counts across cells. Then, we took the natural log of the UMI counts.

### Dimension reduction and visualization of scRNA-seq data

Seurat [62] is used for data integration, data normalization, dimension reduction, cell clustering and other basic scRNA-seq data analyses following our previous studies [37, 63]. To avoid highly expressed genes dominate in later analyses, we scaled the mean and variance of each gene across cells is 0 and 1, respectively. The high variable genes were selected for each scRNA-seq data and PCA was conducted on the scaled highly variable genes. t-distributed stochastic neighbor embedding (t-SNE) is widely used for visualization of scRNA-seq data due to its advantage in showing cell clusters[64]. Unsupervised clustering of the cells was performed using graph-based clustering based on SNN-Cliq [65] and PhenoGraph [66]. We displayed cluster specific expression genes on t-SNE, which provide nice visualization for distinguishing different cell clusters.

### Identification of cluster specific genes and differentially expressed genes

Gene expressions of each investigated cluster were compared to that of remaining clusters by likelihood-ratio test. The genes that are significantly high expressed in the investigated cluster were called as cluster specific genes. The significantly differential expressed genes between two clusters are also tested by likelihood-ratio test. We use Metascape (http://metascape.org) [67] to perform biological process enrichment analysis.

### Gene scoring

To compare gene signatures between sub-populations, we utilized individual gene scores as described previously [68]. Briefly, given an input set of genes (G_i_), we define a score GS_i_(j), for cell j, as the average expression of the gene in G_i_. We also defined a control gene-set based on the expression of input genes. All analyzed genes are binned into 30 bins of equal size, we randomly choose 100 genes from the expression bin of each gene from input genes as control gene-set. We calculated the Z-score of the input genes expression as the final gene set score.

### MetaNeighbor anaylysis

To compare the similarity of cell identities from MSCs with different sources, we performed MetaNeighbor [69] analysis using the R function “run_MetaNeighbor”.

### Single-cell regulatory network inference using SCENIC

We performed SCENIC [70] by starting from the raw counts and following the proposed workflow using the default parameters.

### Statistical analysis

All statistical analyses and graphics were conducted using R. The likelihood-ratio test in Seurat was performed to identify the significantly differential expressed genes between two cell clusters. Bonferroni correction was conducted for multiple testing corrections.

## Acknowledgements

This study was supported by National Key R&D Program of China (2018YFC1004500), National Natural Science Foundation of China (81872330, 81972116), the Shenzhen Innovation Committee of Science and Technology (JCYJ20170817111841427, ZDSYS20200811144002008), the Shenzhen Science and Technology Program (JCYJ20170817172023838, JCYJ20170413161649437), and Center for Computational Science and Engineering, Southern University of Science and Technology.

## Author’s contributions

W.J. conceived the project. X.L. and X.X. collected the samples, cultured MSCs and conducted the chondrogenesis induction. W.H., and C.H., analyzed the scRNA-seq data and conducted comparison analyses, with contribution from P.Q. and N.H.. W.J., L.D., and D.W. supervised this project and interpreted the results. W.H., and W.J. wrote the manuscript, with input from other authors. All authors have read and approved the manuscript.

## Availability of sequencing data

The raw sequence data reported in this paper have been deposited in the Genome Sequence Archive in BIG Data Center, under accession numbers HRA000086.

---

**Figure S1.**
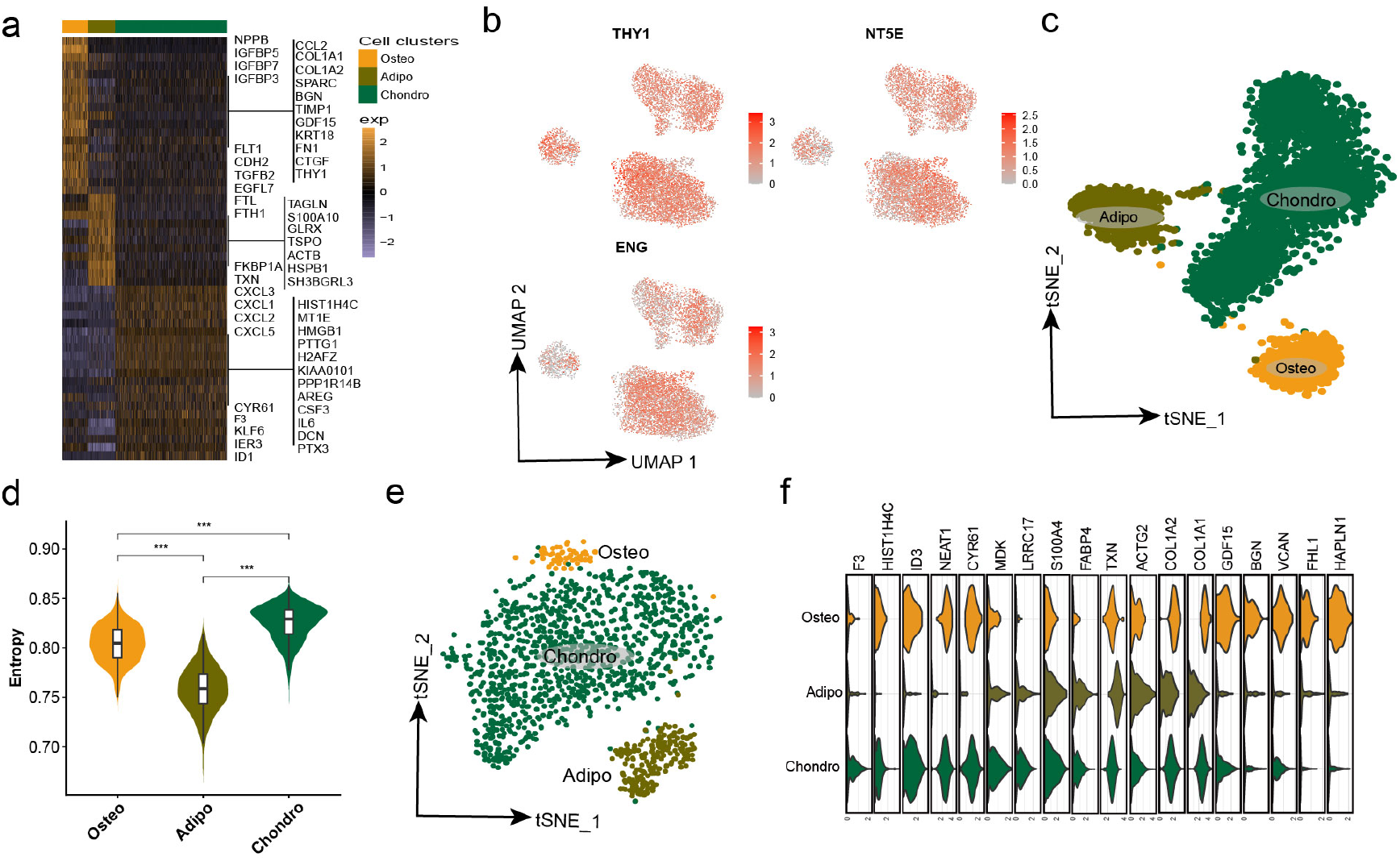
Single cell RNA-sequencing of MSCs and heterogeneity of MSCs. Related to Figure1. a. Heatmap of DEGs in UC-MSCs. b. Feature plots of three marker genes of pooled MSCs from UC, BM, SY and AD. c. t-SNE plot for UC-MSCs showing three typical lineage-specific cell types. d. Violin plot of cellular entropy for each cluster in UC-MSCs. ***: wilcox test, p < 0.0001 e, f. t-SNE plot(e) and typical genes(f) of the validation UC-MSCs.

**Figure S2.**
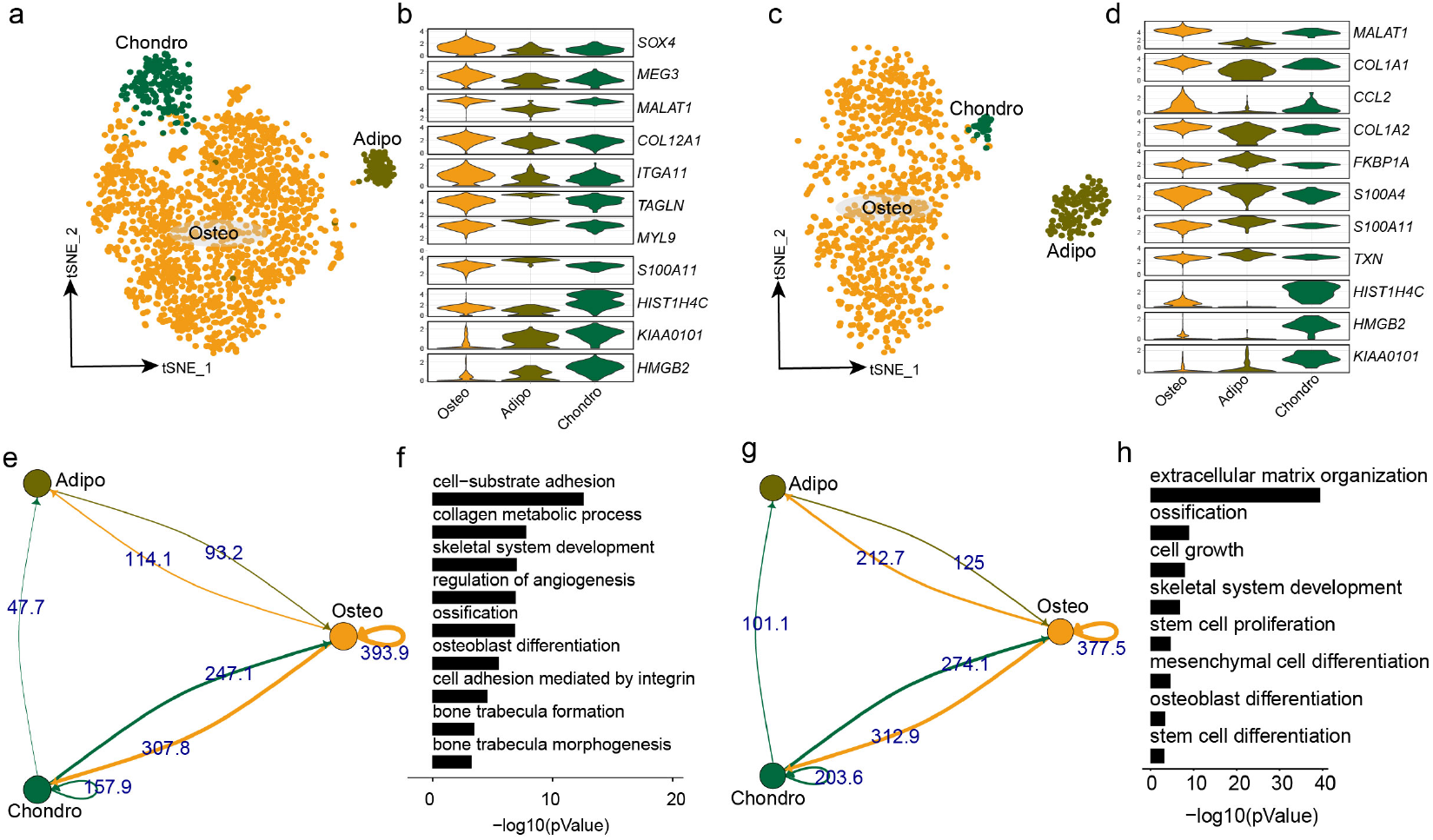
Crosstalk of ligand–receptor pairs among MSCs subpopulations. a,b. Cell identities for BM-MSC (a) and typical genes (b) c,d. Cell identities of SY-MSC (c) and typical genes (d) e. Cell-Cell communication of the pooled MSC samples from UC, BM, SY and AD. f. Highly expressed ligands and receptors (logFC>0.25) of Osteo group in the integrated samples derived enriched GO terms. g. Cell-Cell communication of three cell identities in UC-MSC sample h. Highly expressed ligands and receptors (logFC>0.25) of Osteo group in UC-MSC derived enriched GO terms.

**Figure S3.**
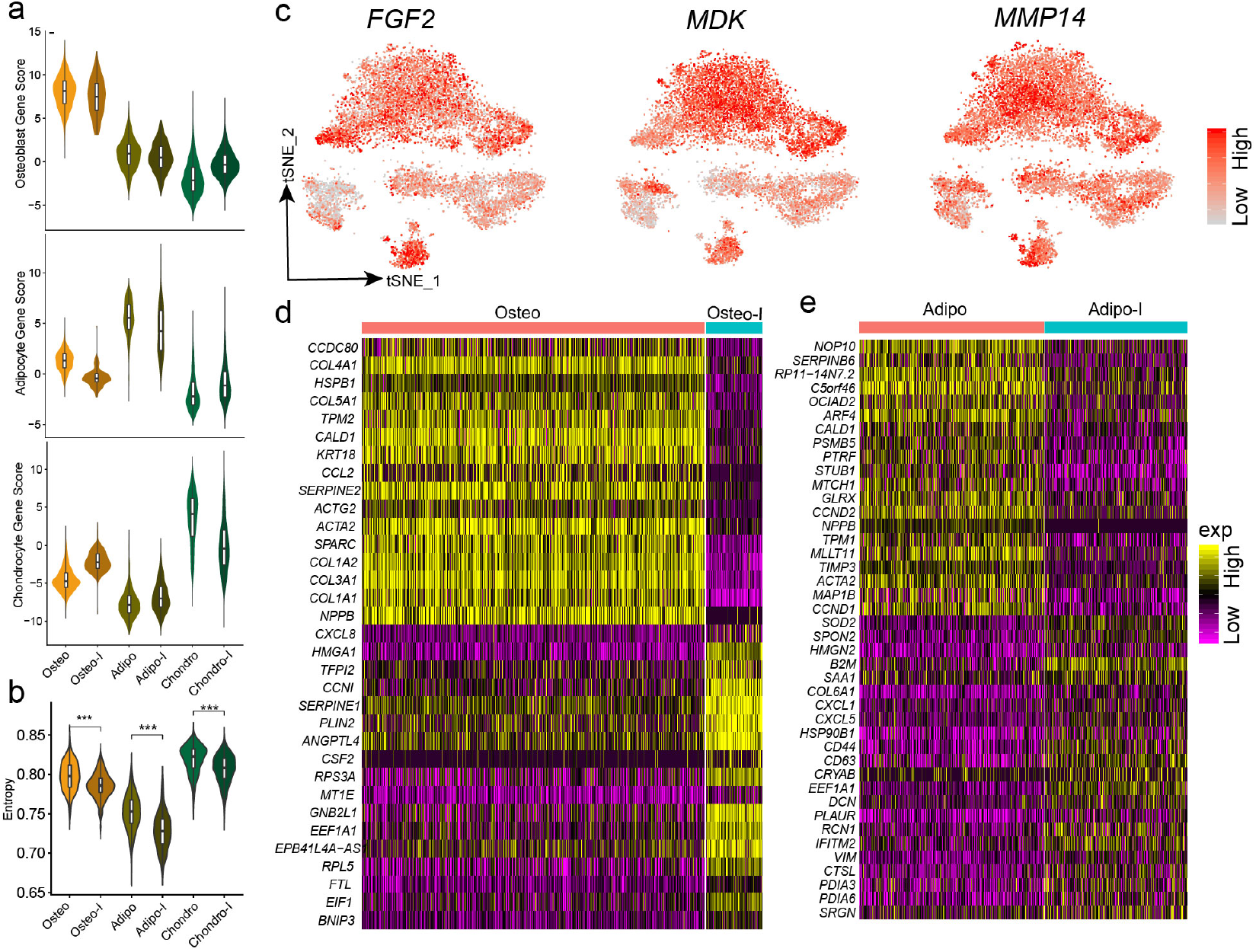
Different cell populations response differentially to chondrogenesis induction. a. Gene score of three cell identities in the integrated UC-MSC and Chondrogenesis Induced UC-MSC. b. Violin plot of cell entropy; ***: wilcox test, p_adj<0.0001 c. Three highly expressed genes in chondrogenesis induced sample than in UC-MSC. d,e. Heatmap of DEGs between the naive and induced sample (Osteo and Adipo).

**Figure S4.**
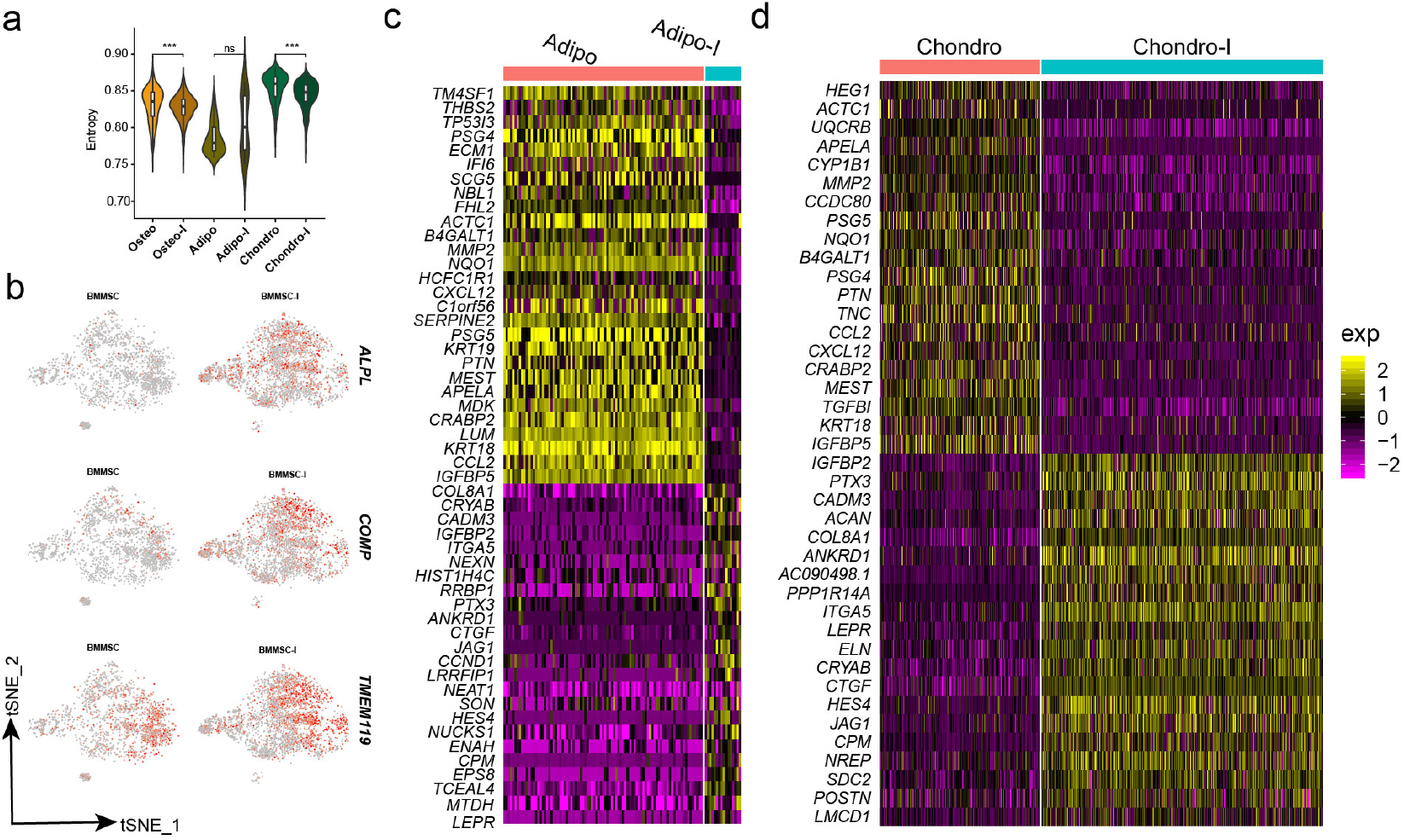
Successful osteogenesis induction in BM-MSC. a. Violin plot of cell entropy; ***: wilcox test, p_adj<0.0001 b. Feature plot showing elevated expression of three genes in osteogenic induced BM-MSC c,d. Heatmaps of DEGs for Adipo-progenitor and chondroprogenitor between the induced and naive samples.

**Figure S5.**
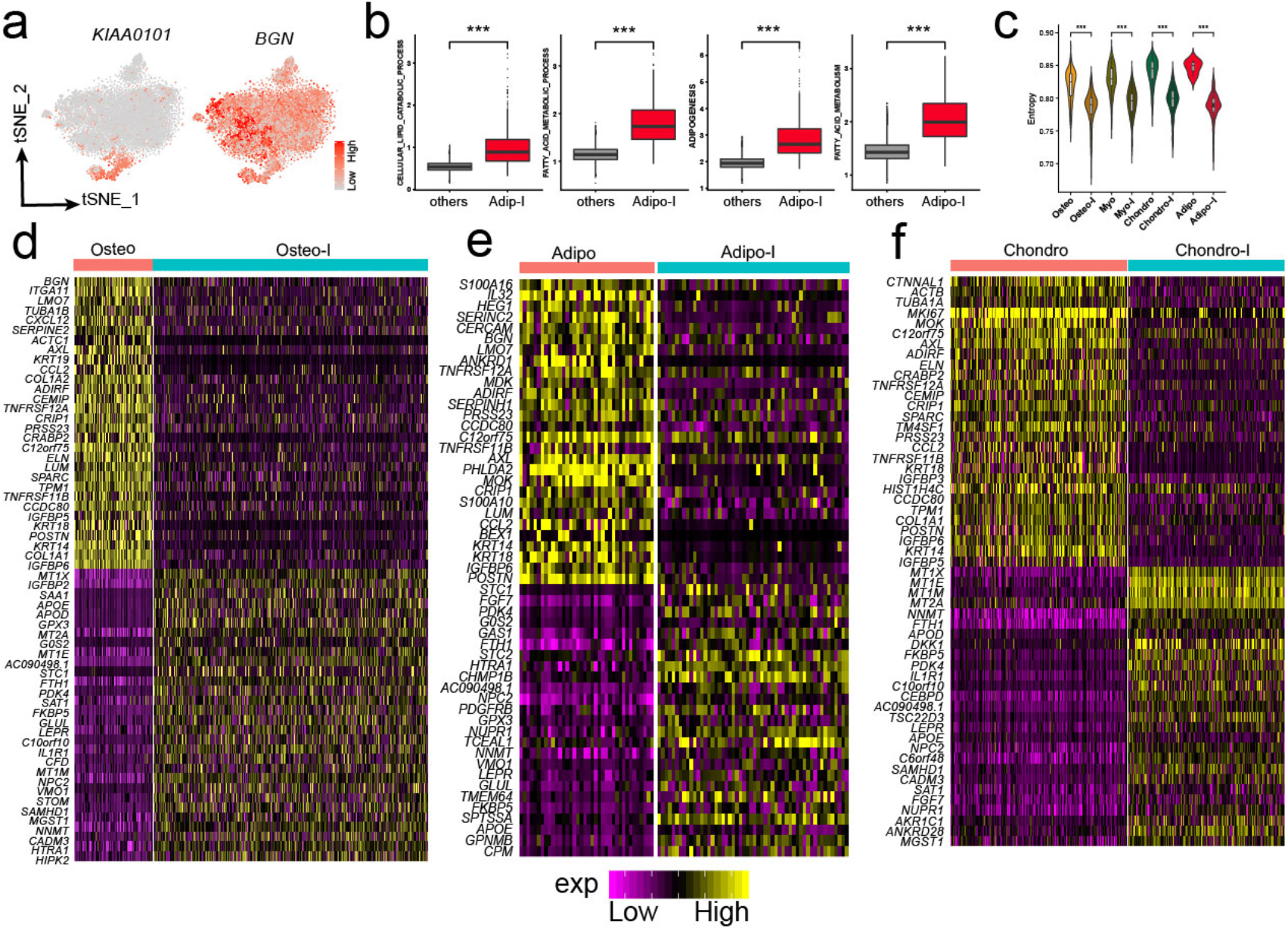
MSC subpopulations response to adipogenesis induction. a. Feature plot of chondroprogenitor and osteoprogenitor represented genes. b. Adipogenesis related pathways significantly enhanced in the adipogenesis induced sample. ***: wilcox test, p < 0.0001 c. Violin plot of cell entropy; ***: wilcox test, p_adj<0.0001 d,e,f. Heatmaps of DEGs for osteoprogenitor, adipo-progenitor and chondroprogenitor between the induced and naive samples.

